# Machine learning driven image segmentation and shape clustering of algal microscopic images obtained from various water types

**DOI:** 10.1101/2024.04.13.589342

**Authors:** Filippo Nelli, Zongyuan Ge, Linda Blackall, Negar Taheriashtiani, Rebekah Henry, Douglas R. Brumley, Michael Grace, Aaron Jex, Michael Burch, Tsair-Fuh Lin, Cheryl Bertelkamp, Anusuya Willis, Li Gao, Jonathan Schmidt, Nicholas D. Crosbie, Arash Zamyadi

## Abstract

Algae and cyanobacteria are microorganisms found in almost all fresh and marine waters, where they can pose environmental and public health risks when they grow excessively and produce blooms. Accurate identification and quantification of these microorganisms are vital for ecological research, water quality monitoring, and public health safety. However, traditional methods of manually counting and morphologically identifying these microorganisms are time-consuming and prone to human error. Application of the machine learning-driven Fast Segment Anything Model (FastSAM), an image segmentation model, automates and potentially enhances the accuracy and efficiency of cell identification and enumeration from microscopic images. We assessed FastSAM for algal cell image segmentation, and three clustering evaluation metrics. Segmentation of microscopic images of algal and cyanobacterial cells in water and treated wastewater samples using the Convolutional Neural Network based FastSAM algorithm demonstrated benefits and challenges of this machine learning-driven image processing. Notably, the pre-trained algorithm segmented entire elements in all microscopic images used in this study. Depending on the shape, 50-100% similarity was observed between machine-based segmentation and manual validation of all segmented elements, with 100% of single cells being correctly segmented by FastSAM. The performance of clustering metrics varied between 57-94% with the Spectral Angle Mapper achieving the most accurate performance, 84-94%, compared to the manually chosen clustering benchmarks. Cyanobacterial and algal communities are biologically diverse and have ecological significance. The application of image clustering techniques in studying their cell shapes marks an important advancement in microbial ecology and environmental monitoring. As technology progresses, these methods will become increasingly utilised to decipher the complex roles that algae and cyanobacteria play in our ecosystems supporting mitigation and public health protection measures.

## Introduction

Algae and cyanobacteria are microorganisms found in aquatic environments worldwide. Cyanobacteria, also known as blue-green algae, are among the most ancient and versatile organisms on earth, playing a pivotal role in various ecosystems (Australian Academy of Science, 2019; Chorus and Welker, 2021). However, the rapid increase in global algal blooms poses significant risks to ecosystems and human health (Water Research Foundation, 2023). Cyanobacterial blooms, often triggered by excess nutrients from agricultural runoff and climate change, can deplete oxygen levels in water bodies, causing dead zones where aquatic life cannot survive (Chorus and Welker, 2021) and this phenomenon has led to massive fish kills in the Darling River in Australia in 2018-19 (Australian Academy of Science, 2019) and in 2023 (Williams and Schulz, 2023).

Furthermore, many cyanobacteria produce cyanotoxins that are harmful to wildlife, livestock, and humans, leading to issues like fish kills, contaminated drinking water, and health problems in humans such as liver damage and neurological effects (Chorus and Welker, 2021; Kibuye et al., 2021a; Kibuye et al., 2021b; Kibuye et al., 2021c; Water Research Foundation, 2023). These toxins include cyclic peptides (microcystins and nodularins), alkaloids (anatoxin-a, anatoxin-a(S), cylindrospermopsins, saxitoxins, lyngbyatoxin-a), lipopolysaccharides, polyketides (aplysiatoxin and debromoaplysiatoxin) and most recently the amino acid β-methylamino-L-alanine (Chen et al., 2017; Shaw et al., 2000; Main et al., 2018; Haddad et al., 2019; Huang and Zimba, 2019; Huang and Zimba, 2019; Nijoy et al., 2019; Zamyadi et al., 2019; Bishop and Murch, 2020; Chorus and Welker, 2021). Non-toxic taste and odour (T&O) compounds produced by algae and cyanobacteria, are the major source of drinking water customer complaints and are difficult to remove from the water (Watson and Jüttner, 2019; Devi et al., 2021; Zamyadi et al., 2021).

Understanding and managing harmful and nuisance algal blooms to mitigate their adverse effects on ecosystems and public health requires detecting the bloom forming microbial cells (Almuhtaram, Kibuye, et al., 2021; Wert et al., 2023). Water utilities across the world need real-time or near real-time monitoring and forecasting tools to implement their mitigation and treatment processes to prevent cells and their harmful and nuisance metabolites from contaminating manufactured water (Treuer et al., 2021; Zamyadi et al., 2021; Water Research Foundation, 2023). Past failures in removing these toxins have led to human exposure, hospitalisation and mortality (Molica et al., 2005; Testai et al. 2016; Lu et al., 2019; Zamyadi et al., 2019; Chorus and Welker, 2021; Devi et al., 2021; Zamyadi et al., 2021).

A traditional way to identify algae and cyanobacteria is via light microscopy to observe their morphological features (Rippka et al., 1979), and they can also be quantified. However, imaging the diverse algal and cyanobacterial cells for their quantification and identification presents several technical challenges due to their inherent morphological variability and environmental adaptability (Zamyadi et al., 2016; Zamyadi et al., 2019; Chorus and Welker, 2021; Vaughan et al., 2022), and their small (> 20 µM diameter) size (Wetzel and Likens, 1991; Vaughan et al., 2022). Cyanobacteria exhibit at least nine shapes (Allaf and Peerhossaini, 2022), four cell types (vegetative, heterocysts, akinetes and hormogonia) and several sizes. Cyanobacteria have developed different shapes over evolutionary time and environmental conditions (e.g., fresh through marine water, eutrophic or oligotrophic nutrient status, temperature ranges from hot (hot springs) through to cold (polar), light availability and strength, and ambient chemical composition) along with their dynamic interactions with other organismal community members can modulate cell shapes and sizes. The morphological diversity alone makes it challenging to use imaging for identity, but this is even more complicated as the same species might have different morphologies at different growth stages (Zamyadi et al., 2016; Jin et al., 2018; Zamyadi et al., 2019; Springstein et al., 2020; Chorus and Welker, 2021; Vaughan et al., 2022) and different species with the same morphology could co-exist.

The phenotypic feature of cell pigmentation provides another identification tool but also challenge (Zamyadi et al., 2016; Chorus and Welker, 2021; Vaughan et al., 2022). Pigments such as chlorophyll, phycocyanin, and phycoerythrin, give them distinct colours ranging from green to blue-green and even red but this feature can affect their optical properties and interfere with fluorescence measurements (Zamyadi et al., 2016; Chorus and Welker, 2021), thus complicating the use of standard light microscopy techniques, as different wavelengths might be required to optimally image different species or to highlight specific cellular components (Zamyadi et al., 2016; Jin et al., 2018; Zamyadi et al., 2019; Chorus and Welker, 2021; Vaughan et al., 2022). The small size of many cyanobacterial cells requires advanced microscopy techniques, e.g., electron or super-resolution light microscopy, to resolve fine structural details. These techniques are expensive, time-consuming, and often require complex sample preparation that could alter the natural state of the cells (Vaughan et al., 2022).

Several methods for algae and cyanobacteria imaging have been developed over time (Chorus and Welker, 2021; Vaughan et al., 2022). Reviewing strengths and limitations of these methods (Supplementary Information Table SI-1), and their application for research and management of manufactured water have shown these methods are demanding in terms of time and specialized expertise limiting their usage for the general water industry to exploit them for monitoring bloom initiation, development and collapse (Lund et al., 1958; Weber, 1973; Wetzel and Likens 1991; Acker, 2003; American Public Health Association et al. 2017; Chorus and Welker, 2021; Vaughan et al., 2022).

Machine learning (ML) involves training machines to learn from data, identifying patterns, and making decisions with minimal human intervention (Almuhtaram, Zamyadi et al., 2021; Centenaro et al., 2021; Rubbens et al., 2023; Vaughan et al. 2023; Zhang et al., 2023).

Artificial Intelligence (AI) extends this by enabling machines to perform tasks that typically require human intelligence, such as visual perception, and decision-making. AI-driven image analysis is pivotal in scientific research, from studying microscopic cells to exploring astronomical images, analysing satellite and aerial imagery for environmental monitoring, urban planning, and disaster management (Rubbens et al., 2023). It automates the detection of patterns that are often imperceptible to the naked human eye, facilitating ground-breaking discoveries (Centenaro et al., 2021; Rubbens et al., 2023).

These technologies are increasingly becoming integral partners in exploration, discovery, and innovation, and monitoring offering ground-breaking advancements in both the quality and efficiency of image analysis (Almuhtaram et al, 2021; Rubbens et al., 2023; Zhang et al., 2023). A key application has been in image classification, where ML algorithms are trained to identify and classify various biological structures in images, such as cells, tissues, or specific proteins (Williamson et al., 2020; Winfree, 2022; Rubbens et al., 2023; Zhang et al., 2023). In imaging and analysis, ML and AI play crucial roles in diagnosing patterns (Williamson et al., 2020; Winfree, 2022; Rubbens et al., 2023). These algorithms can detect abnormalities with high precision, leading to early and more accurate diagnoses, improving intervention outcomes (Wang et al., 2022; Winfree, 2022). Expanding these applications to algae and cyanobacteria monitoring is challenging considering difficulties in algal and cyanobacterial imaging, including variability in shape, size, and colour. However, this has the potential to provide cost effective accurate diagnostic tool for precise real-time identification of cells and on-time implementation of management strategies.

In the field of digital imaging including digital microscope photos, AI technologies like neural networks, particularly Generative Adversarial Networks (GANs), have revolutionized image synthesis, enabling the creation of high-quality, lifelike images from scratch (Rubbens et al., 2023). Convolutional Neural Network (CNN) is a type of deep learning algorithm particularly well-suited for analysing visual imagery. CNNs are widely used in various fields, including biology and medicine, for tasks like cell classification, disease detection, and medical image analysis (Rubbens et al., 2023; Vaughan et al., 2022). In cell imagery, CNNs and CNN based algorithms such as the Segment Anything Model, can be employed and fine-tuned to recognize various types of cells, detect abnormalities, or even track cell movements in time-lapse images. For example, they can be used to distinguish between healthy cells and damaged cells, or to identify other cellular structures. The strength of CNNs in cell imagery lies in their ability to learn features directly from the image data with minimal pre-processing and labelling, capturing intricate patterns that might be missed by human observers or traditional computer vision techniques (Rubbens et al., 2023; Vaughan et al., 2022).

Machine learning has also enabled automated image segmentation, which involves separating different features within an image for detailed analysis. This is particularly useful in quantitative cell imaging, where it’s important to distinguish individual cells or subcellular structures for further study (Rubbens et al., 2023). The Fast Segment Anything Model (FastSAM) algorithm leverages the computational prowess of CNNs to deliver solutions in real-time for the ‘segment anything’ task (Kirillov et al. 2023; Zhao et al. (2023).

Integration of machine learning with imaging has facilitated large-scale analysis of biological data, allowing researchers to process and interpret vast amounts of imaging data far more rapidly and accurately than traditional methods (Keller et al., 2018; Wacquet et al., 2018; Rubbens et al., 2023). This scalability is crucial for high-throughput studies, such as in drug discovery or genomics, where large datasets are common. Machine learning in biological imaging has not only improved the accuracy and efficiency of image analysis but also opened new avenues for research and discovery in the biological sciences (Keller et al., 2018; Rubbens et al., 2023). Advancements in these areas are crucial to improve the reliability and utility of machine learning applications in algae and cyanobacteria imaging.

Hence, this study aims to explore the capabilities of machine learning algorithms to segment and cluster algae and cyanobacteria cells from freshwater reservoirs and treated wastewater stabilisation ponds. The specific objectives are (i) to segment cells and cell formations considering background water quality which impacts quality of the image, (ii) to design an unsupervised clustering approach to group cells following the ground truth, and (iii) to assess the performance of the segmentation and clustering algorithms and conduct a quality control check. To the best of the authors’ knowledge, this paper presents novel results exploring the usage of AI-based segmentation algorithms coupled with automated clustering approaches modified for real world samples from water bodies and treated wastewater stabilization ponds with varying water quality backgrounds. This study focuses on image segmentation and clustering as the key steps prior to identification of the clustered groups which is subject of the authorship team’s ongoing research work.

## Material and Methods

### Data Collection

For the purpose of this study, a meticulously curated dataset comprising 1,007 colour microscopic images of cyanobacteria and algae within both freshwater sources and treated wastewater stabilization ponds in Victoria (Australia), was assembled. This collection was designed to encapsulate a broad spectrum of cell morphologies, sizes, and ecological contexts. The procurement of these images was exclusively facilitated through the utilization of phase contrast microscopy techniques.

The dataset encompasses images captured via two distinct models of microscopy cameras: a total of 685 images were acquired using a ZEISS Axiocam ERc 5s camera, whereas a smaller subset of five images was obtained with an Olympus EP50 camera. Additionally, a subset of 317 images was incorporated into the dataset without specific camera metadata. The magnification levels at which these images were captured varied, with 92 images recorded at 100× magnification, 484 images at 200× magnification, and the remaining 431 images at 400× magnification. It is noteworthy that all images were processed while retaining their original magnification settings.

To enhance the analytical utility and uniformity of the image dataset, a series of preprocessing techniques were meticulously applied using scikit-image Python library. These included: (i) the application of noise reduction algorithms to sharpen edge definition, thereby facilitating more precise object segmentation (Python setting: sigma_color = 0.05, sigma_spatial = 15, patch_size = 5, patch_distance = 3, h = 0.1); (ii) contrast enhancement to augment the visual differentiation of features within each image (Python setting: clip_limit = 0.03); and (iii) color normalization procedures aimed at optimizing the efficacy of the mask generation phase of the segmentation methodology.

### Machine learning workflow

The framework for segmentation and clustering was devised with the objective of evaluating the precision with which the implemented algorithms can segment and categorize cyanobacterial and algal cells within the digital images. To facilitate a comprehensive understanding of this process, a workflow diagram alongside corresponding pseudocode is delineated in Figure 1. This illustration serves to provide a succinct graphical representation of the sequential steps undertaken within the methodological approach, highlighting the critical pathways through which the algorithms process and analyze the digital images to achieve cell segmentation and grouping.

**Figure 1.**
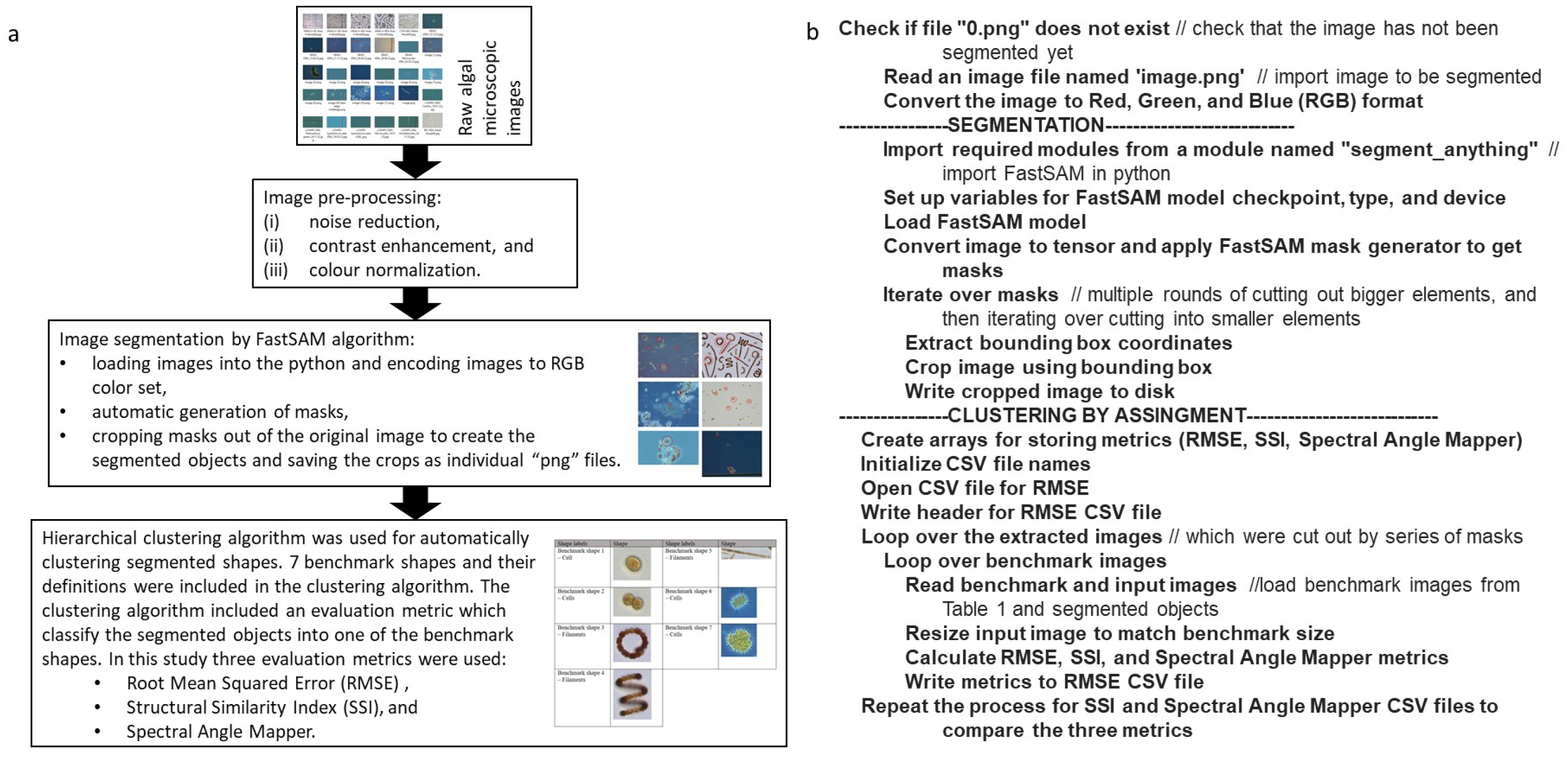
The segmentation and clustering process (a) workflow, and (b) pseudocode.

### Segmentation Algorithms

In the context of this research, the delineation of segmentation categories was informed by a combination of empirical observations of algal blooms within the study sites and established definitions provided by the Australian Academy of Science (2019) and Chorus and Welker (2021). The operator-defined categories for segmentation purposes include “Single cell”, “Filaments”, “Cells out of filaments”, “Colonies”, and “Cells out of colonies”. These classifications facilitate a structured segmentation of cyanobacterial and algal populations by categorizing the diverse morphologies encountered in aquatic environments.

The analytical foundation of this study is anchored on the FastSAM algorithm, an approach to image segmentation that leverages a CNN for real-time processing. This algorithm, adapted from the methodologies proposed by Kirillov et al. (2023) and Zhao et al. (2023), is suitable for the segmentation of cyanobacterial and algal cells. The FastSAM algorithm incorporates a set of pre-selected parameters including light tolerance, brightness, and contrast, which are dynamically adjusted by the algorithm to refine its performance. This fine-tuning process is crucial for enhancing the algorithm’s efficacy in processing the unique visual characteristics of cyanobacteria and algae imagery.

FastSAM manages to significantly cut down on computational and resource demands while maintaining a performance level that is on par with its predecessor, SAM, thus facilitating its application in real-time scenarios (Kirillov et al., 2023; Zhao et al., 2023). At the core of FastSAM’s effectiveness is the YOLOv8-seg object detection architecture, which incorporates an instance segmentation branch, thereby empowering the algorithm to accurately generate segmentation masks for all discernible objects within an image.

For the purposes of this study, the FastSAM algorithm was operationalized through a series of methodically structured implementation steps, as follows:

1. Image Importation: Digital images were imported into the Python programming environment, adhering to a resolution benchmark of 1920×1080 pixels.
2. Color Space Conversion: These images were then converted into the Red, Green, and Blue (RGB) color spectrum to facilitate accurate color representation and analysis.
3. Automated Mask Generation: Utilizing the open-source database of 1 billion Synthetic Aperture Masking entries, FastSAM generated masks by setting the parameters to process 8 points per batch with a confidence threshold of 0.6. This process also included the generation of additional crops from segmented objects, leveraging the algorithm’s capability to initiate further detailed segmentations from already segmented objects.
4. Mask Extraction and Object Segmentation: Masks were extracted from the original images, which enabled the production of precisely segmented objects.
5. Storage of Segmented Outputs: The resulting cropped segments were stored as individual Portable Network Graphics (png) files, with dimensions varying according to the segmentation details.

The term “points per batch” refers to the quantity of points assessed against existing masks for each object, with a set value of 8 points per specific batch in this instance. This assessment aids in identifying any correspondence with existing masks, where a higher point count improves accuracy but demands more memory. The “internal confidence threshold” is defined as the minimum confidence value, in this case 0.6, required for masks to be deemed valid, with higher thresholds imposing stricter validation criteria. To evaluate the accuracy of machine-based segmentation against predefined categories, a manual review process was implemented.

The image pre-processing and segmentation algorithm were executed on a High-Performance Computer (HPC) utilizing 512 cores in parallel, which resulted in a 50% reduction in computational time, underscoring the algorithm’s efficiency and suitability for large-scale, real-time applications (allowing 8 points per batch with a confidence threshold of 0.6).

### Shape Clustering Techniques

In the preparation phase of this study, the utilized images underwent a process of manual annotation, guided by insights from experts in microbiology. The objective was to identify seven distinct clustering benchmark shapes, as delineated in Table 1. This annotated image set was employed to assess the performance of the clustering algorithms. To uphold the integrity of the study, thorough quality control measures were instituted, aiming to verify both the precision and the consistency of the annotations applied to the images. The selection of these specific shapes, as itemized in Table 1, was intentional and served a dual purpose. Primarily, it was designed to anchor this proof-of-concept investigation, which concentrates on the delineation and analysis of individual cells and filaments. Secondly, it facilitated a focused evaluation of the clustering algorithms’ capability to categorize these biological entities based on their morphological characteristics.

**Table 1.**
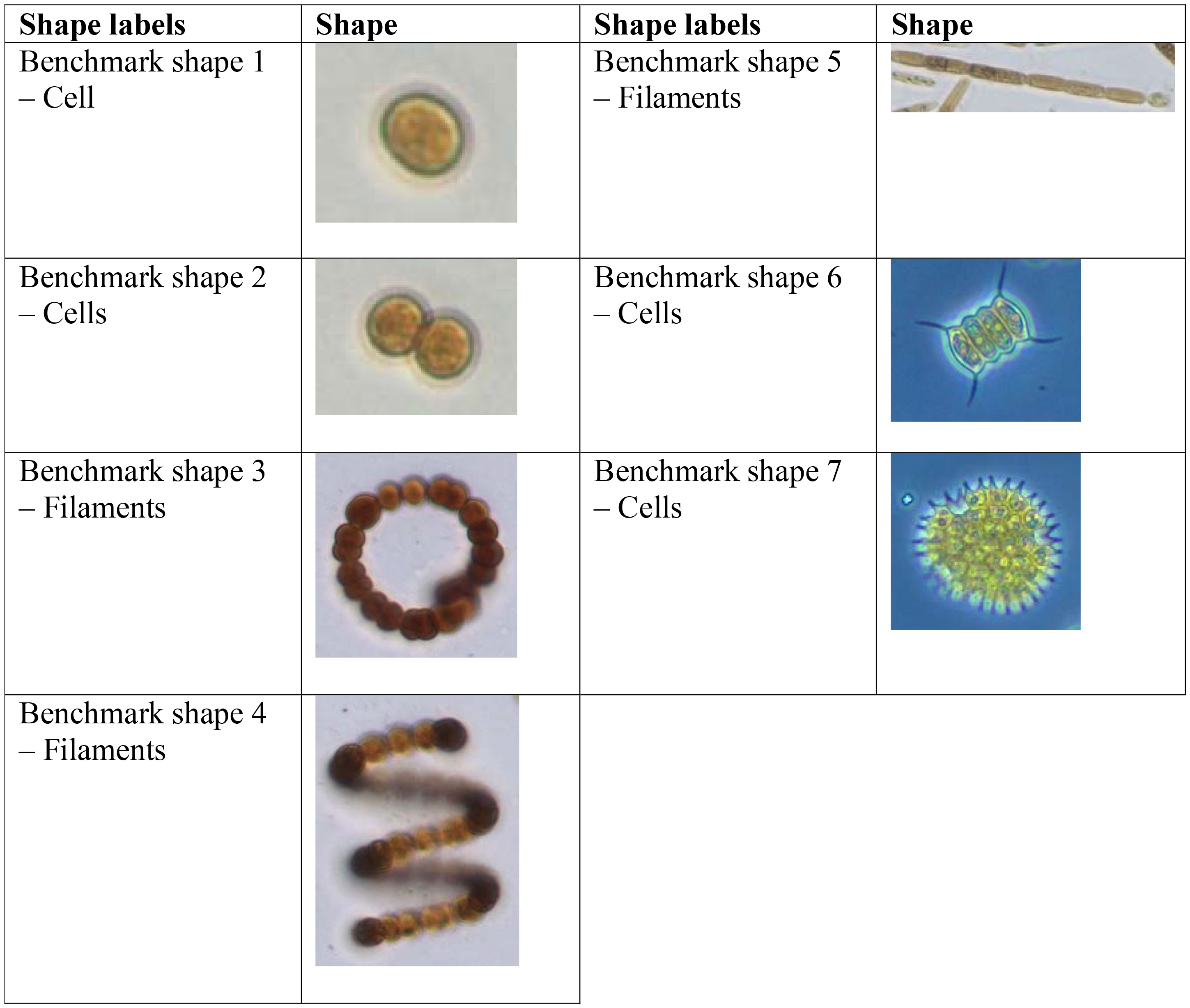
Seven manually chosen benchmark categories for clustering purposes.

| Shape labels | Shape | Shape labels | Shape |
| --- | --- | --- | --- |
| Benchmark shape 1<br>– Cell |  | Benchmark shape 5<br>– Filaments |  |
| Benchmark shape 2<br>– Cells |  | Benchmark shape 6<br>– Cells |  |
| Benchmark shape 3<br>– Filaments |  | Benchmark shape 7<br>– Cells |  |
| Benchmark shape 4<br>– Filaments |  |  |  |

Benchmark clustering algorithm, which is clustering by assignment (Figure 1b), was used for automatically clustering segmented shapes. This algorithm can detect subtle differences in shapes and texture that might be overlooked by the human eye. The clustering algorithm included an evaluation metric which classify the segmented objects into one of the 7 benchmark shapes. If an object didn’t match the benchmark criteria then it was flagged and separated into “unknown” category. In this study three evaluation metrics were tested by running the clustering algorithm separately for each metric.

Several evaluation metrics are available. Developing robust evaluation metrics for the clustering process was a crucial step in this study. Effectiveness of clustering methods heavily relies on the precision and reliability of the evaluation metrics used. This study involved adaptation of three evaluation metrics, calibration, and statistical analysis to assess the performance of the clustering metrics. Calibration processes were undertaken by establishing metric-specific thresholds that ensured an optimal correlation between segmented objects and benchmark shapes. These thresholds were informed by the findings of a prior study (Nelli et al., 2023), adopting a consistent mathematical framework for analysis. Clustering metrics included the evaluation of similarity between clustering outcomes and benchmark shapes (Table 1), using a Python-based similarity algorithm. This algorithm, leveraging the *sewar* library which is a python package, incorporates the following metrics (Figure 1a): (i) Root Mean Squared Error (RMSE), which quantifies the alignment between segmented objects and benchmark thresholds; (ii) Structural Similarity Index (SSI), which evaluates similarities in image structure (Wang et al., 2004); and (iii) Spectral Angle Mapper, which aligns image spectra with known spectra (Rashmi et al., 2014).

To ensure the precision of the clustering methodology, manual quality-control assessments were conducted to compare the accuracy of AI-driven classification against the benchmark shapes (Table 1). Statistical analyses, incorporating ANOVA and t-tests, were executed to determine the statistical significance of the outcomes. Moreover, confidence intervals for the metric scores were calculated, offering insights into the variability and reliability of the algorithmic performance across different evaluations.

## Results and Discussion

Figure 2 presents specific examples of findings of this study on the effectiveness of used CNN-based segmentation algorithm. Segmentation is the first key step in digital image processing, particularly in the analysis of biological images. This segmentation process enabled automated isolation of pre-determined categories, which is essential for understanding the growth patterns, behavior, and interactions of microorganisms like cyanobacteria. Further examples of image segmentation to show the diversity of cells and colonies that the algorithm was able to successfully separate are presented in Supplementary Information Table SI-2. These images provide additional visual evidence of the versatility of the process used in this study.

**Figure 2.**
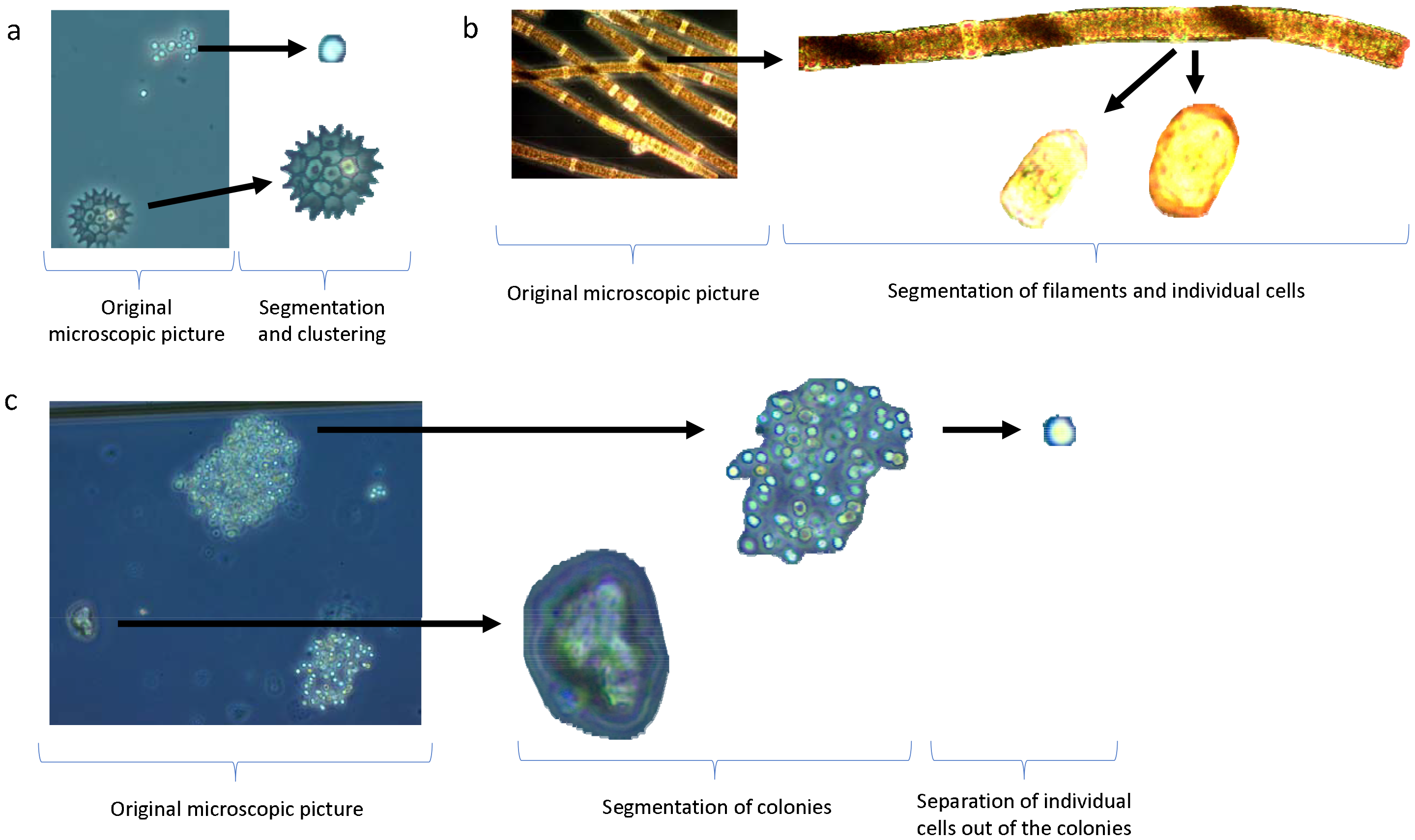
Image segmentation and clustering: (a) simple but mixed real-world sample; (b) uni-species real world filamentous sample; (c) complex real-world samples with mixed species and agglomerations (colonies).

Figure 2a demonstrates the process of image analysis where the original microscopic image has been subjected to the segmentation algorithm. The right side of the image shows the result after segmentation has been applied. The arrows point to specific structures within the original image, and the corresponding segmented representations of those structures. The segmentation process has partitioned the digital image into multiple segments to make the image more meaningful and easier to analyze. These segmented images will then be clustered,grouping the set of objects in such a way that objects in the same group are more similar to each other than to those in other groups.

Figure 2b is a visual representation of the segmentation process applied to filamentous cyanobacteria. The image on the left shows a filamentous structure of cyanobacteria under a microscope, characterized by the chains of cells. In the middle part of the image, an arrow is pointing to a processed version of the filamentous structure where the segmentation has been applied. This segmentation process involved differentiating the individual cells from the background and from each other, a crucial step for quantitative image analysis. On the right of Figure 2b, there are two individual cells that have been segmented from the filament. The arrows point to these isolated cells, highlighting the results of the segmentation process. This level of detail would be useful for analyzing characteristics of individual cells, such as their size, shape, and perhaps the internal structures visible within the cells.

CNN-based algorithms automatically and adaptively learn spatial hierarchies of features from input images. This is crucial in cell imagery where the visual patterns, such as the shape and size of cells, are important for classification or analysis (Vaughan et al., 2022; Kirillov et al., 2023; Rubbens et al., 2023). Convolutional Layers are the core building blocks of a CNN.

They apply a convolution operation to the input, passing the result to the next layer. This process involves filtering the image with smaller, learnable filters, allowing the network to capture important features like edges, textures, and other cellular characteristics. Pooling layers reduce the spatial size of the convolved features, helping in decreasing the computational power required to process the data (Kim et al., 2023). They also help in making the detection of features invariant to scale and orientation changes. After several convolutional and pooling layers, the high-level reasoning in the neural network is done through fully connected layers. In cell imagery, this part of the network would interpret the features extracted by the convolutional layers to classify different cell types or identify anomalies.

The segmentation of 1007 diverse microscopic images of water samples collected from the field, using the CNN-based algorithm, represents a significant advancement in the field of image processing and biological analysis. A maximum of 41 distinct shapes were segmented in one sample. Table 2 shows FastSAM was able to identify individual cells, colonies, and filaments out of every picture without issues. The algorithm was able to achieve 50-100% correct segmentation of elements in each picture for each category (Table 2). However, the segmentation ability declines whenever smaller elements need to be extracted out of larger ones. In this case the model suffers the lack of contrast between the object, which are always surrounded by copies of themselves. In the most complicated cases, the algorithm was unable to separate the individual elements, limiting the segmentation analysis to smaller part of the colonies.

**Table 2.** Percentage of elements correctly segmented in each picture for each category. Data represents the comparison between the machine-based segmentation and human-based segmentation.

| Segmentation categories | % of correct segmentation |
| --- | --- |
| Single cell | 99.6% |
| Filaments | 99.7% |
| Cells out of filaments | 49.6% |
| Colonies | 99.5% |
| Cells out of colonies | 60.2% |

CNNs, known for their efficacy in image recognition and classification tasks, were adapted here for a more nuanced and specialized task of microscopic segmentation. The primary achievement of this study lies in the CNN’s ability to distinguish between individual cyanobacteria cells within a clustered and visually complex environment. This task involved overcoming challenges such as varying cell shapes, overlapping cells, and inconsistent lighting conditions in microscopic imagery. Achieving a balance between sensitivity (detecting as many true cells as possible) and specificity (avoiding false positives) was critical in such segmentation tasks. Notably, false positives are segmented objects that are not actual objects so the non-relevant portion of an image.

Other studies, such as Jo et al. (2017), have employed similar CNN-based approaches for segmenting microorganisms in water samples. To improve their degree of accuracy they have used an advanced quantitative phase imaging unit and were able to achieve up to 96.3% recognition accuracy (Jo et al., 2017). However, the degree of accuracy in segmenting densely populated images under microscope as achieved in the current study for small sized cyanobacterial cells is unique. Traditional methods for cyanobacteria segmentation, like manual counting or basic image processing techniques, often fall short in terms of accuracy and efficiency (Vaughan et al., 2022). The CNN-based approach in this study demonstrates a significant improvement over these traditional methods. Previous studies have explored different CNN architectures within a multiclass arrangement where each class represented a particular species of cyanobacteria (Jo et al., 2017; Krizhevsky et al., 2017; Sun et al., 2018; Wahid et al., 2018; Shashni et al., 2018; Kwabena Patrick et al., 2019). It was concluded that performance of recognition algorithms is affected considerably by these arrangements (Jo et al., 2017; Krizhevsky et al., 2017; Sun et al., 2018; Wahid et al., 2018; Shashni et al., 2018; Kwabena Patrick et al., 2019; Rubbens et al., 2023; Vaughan et al., 2022).

In a previous study (on manual labelling and R-CNN) cyanobacterial and algal cells in the images were manually labelled with their scientific names and positions to train Fast R-CNN models (Rubbens et al., 2023). Compared to that study Figure 2b segmentation correctly identifies the presence of colonies and filaments, cutting them out from the remaining portion of the image. The pre-trained “1.1 billion segmentation masks (SA-1B dataset)” recognized all category shapes in all microscopic images used in this study (Figure 2 and Table 2).

Cao et al. (2022) discussed the segmentation of cyanobacteria images from Taihu Lake using CNN-based algorithms and noted that the outline of cyanobacteria lacked detail and there were many holes in the images. This highlights the challenges in the segmentation process, such as the loss of details due to the algorithm not fully considering the relationship between pixels. The higher quality of microscopy images used in this study reduced the likelihood of segmentation issues. However, while the segmentation algorithm used in this study did not face any difficulty in recognising large colonies and filaments or individual cells (Figure 2), the quality of the outcomes becomes lower for elements extracted out of them depending on the quality of floc image. The main issue in these cases, was the lack of clean edges between cells inside larger elements, which made it hard for the automated algorithm to detect them as separate objects. It is possible to overcome this issue using a multi threshold approach, which would run the segmentation algorithm using higher thresholds (to extract colonies and filaments) and lower thresholds (for less clear elements). More tests are required to identify the optimal values of the segmentation algorithm parameters to avoid detecting non-existent objects such as shades and duplicates.

Segmentation of images used in this study are quite clear and detailed (Table 2), due to the use of advanced segmentation methods. Image pre-processing was the initial step which involved noise reduction, contrast enhancement, and colour space conversion to improve image quality for subsequent analysis. Gaussian blurring and histogram equalization were employed to address issues such as uneven lighting and to enhance image details automatically. A combination of thresholding, edge detection, and region-based segmentation was used to improve the process of dividing images into meaningful regions. The choice of method depended on the specific characteristics of each image, such as contrast between the organisms and the background. Feature extraction involved identifying distinctive attributes of the segmented regions. This step was pivotal in differentiating between various types of shapes. Finally, recognition involved classifying the extracted features into specific categories. The CNN-based algorithm showed promising results in achieving high accuracy in this step.

Clustering results are presented in Figure 3. Current challenges for clustering by assignment include differentiating between cyanobacteria and other microorganisms, dealing with varying shapes and sizes, and adapting to different lighting conditions in images. From the studied clustering criteria, the spectral angle mapper was the best indicator (Figure 3). It was able to identifying and eliminating the segments that do not belong to any of the seven benchmarks, the “unknown” group, in 83-94% of cases, in particular for clear objects such as Benchmarks 6 and 7. RMSE and SSI percentages were significantly lower compared to the spectral angle mapper, highlighting knowledge gaps in the clustering model. The poor outcome of the two indicators was due to the mistake in clustering similar elements together, for example Benchmark 2 instead of Benchmark 1 or Benchmark 6 instead of Benchmark 7. The RMSE in particular was not precise enough in distinguishing between Benchmark 2 similarities, often confusing single cells for colonies and vice versa. Similarly, the SSI indicator struggled to recognise the difference between benchmark 6 and 7, due to the similar chromatic features of the images. However, grouping together benchmark 1-2, 3-4,5,6-7 into four, more different groups provided significantly better results with percentages in line with spectral angle mapper results, showing the ability of the clustering algorithm in correctly recognizing the type of segmented objects. Variations in image quality, overlapping shapes, and the sheer diversity of cyanobacterial forms make algorithmic classification a complex task. Researchers continually refine these algorithms for greater accuracy and efficiency in identifying and categorizing cyanobacterial shapes (Jo et al., 2017; Krizhevsky et al., 2017; Sun et al., 2018; Wahid et al., 2018; Shashni et al., 2018; Kwabena Patrick et al., 2019; Rubbens et al., 2023; Vaughan et al., 2022).

**Figure 3.**
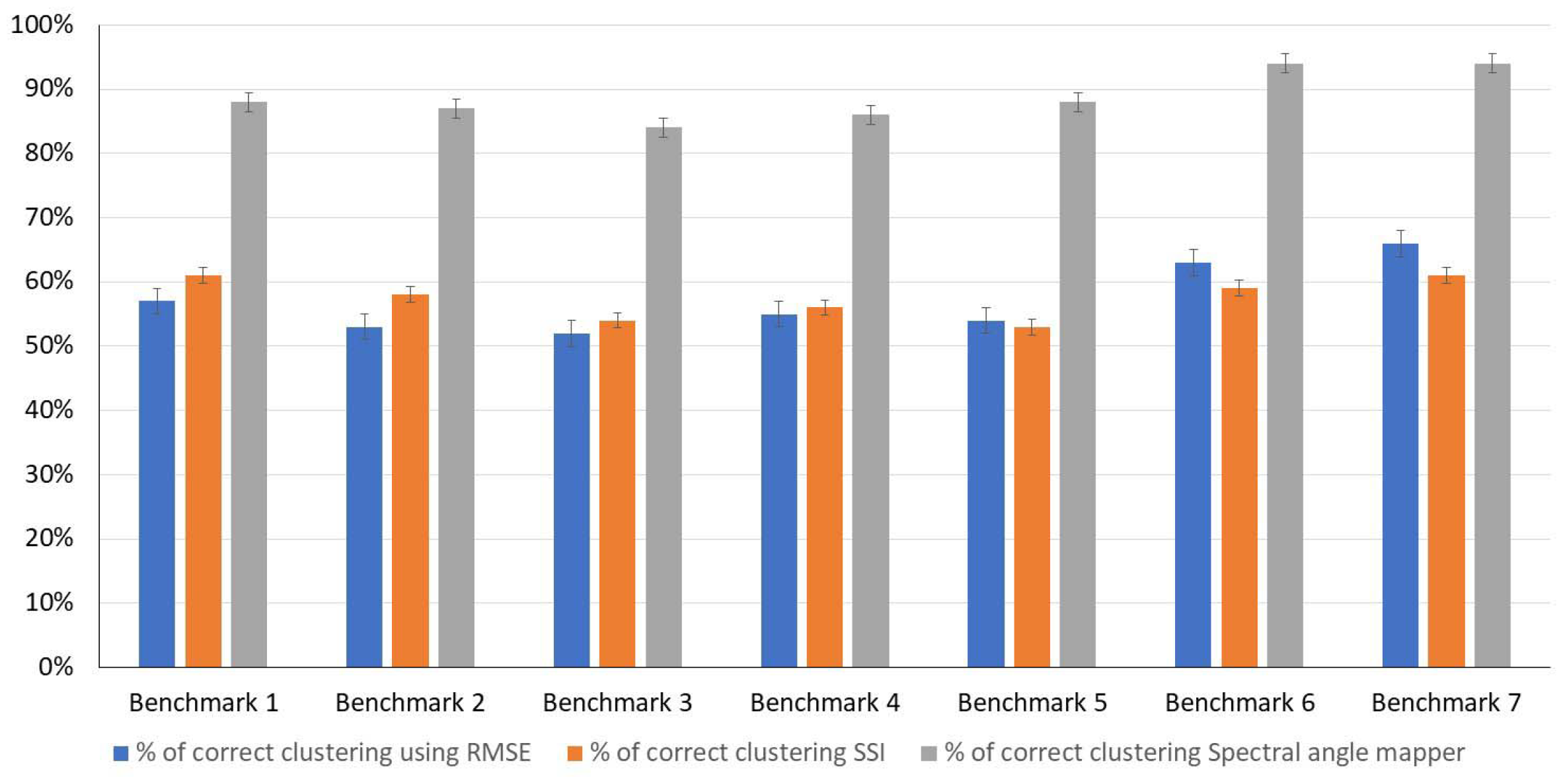
Percentage of correct clustering for each one of the tested evolution metrics and benchmark shapes (as defined in Table 1). Notably, the difference to 100% was classified into the “unknown” group.

While the segmentation algorithm based on Spectral Angle Mapper is able to provide results of good quality already at this stage, the clustering section can only predict the general classification of the microorganism but is unable to predict reliably the total number of them or the specific differences. Additional indicators could be included in the model to boost the analysis of the image structure (For example Universal Quality Image index - UQI) and refine the clustering ability of the code. Another approach worth exploring is the introduction of a second neural network, in addition to Spectral Angle Mapper, to consider the classification of the segments. For example, OpenAI Contrastive Language-Image Pretraining (CLIP) showed promising results in analyzing image similarities (Hentschel et al., 2022) and it could be adapted for this purpose after a proper training phase.

Based on the outcomes of this study further explorations are required to address limitations and challenges related to methodological details and biological applications including segmentation accuracy, adaptability to variations, feature extraction and representation, dataset quality and availability, interpretability, real-time processing capabilities, robustness to imaging conditions, handling noise and artefacts, deep learning techniques, creation of synthetic micrographs, integration with other omics data, with automatic sampling and microscopic imaging systems, with biological knowledge and modelling, and with toxins or T&O compounds production.

Machine learning models may struggle with accurately segmenting cyanobacteria due to their diverse morphologies and overlapping structures in dense populations. This could lead to errors in identifying individual cells or colonies, affecting subsequent analysis. Cyanobacteria and algae exhibit significant variation in shape and size across different species and environmental conditions. Machine learning models may not generalize well across these variations, leading to decreased accuracy in unfamiliar samples. Furthermore, effective feature extraction and representation are crucial for shape clustering. There might be a gap in identifying and extracting relevant features that accurately represent the unique characteristics of cyanobacteria. Clustering algorithms require further tuning using larger number of high-resolution images to enhance the performance of the metrics.

The quality and availability of annotated datasets for training and validating machine learning models are critical. There might be a scarcity of high-quality, diverse datasets representing different cyanobacteria types under various conditions. Many advanced machine learning models, often act as “black boxes.” There is a need for more interpretable models to understand how decisions are made, which is crucial for scientific research.

For practical applications, the ability to process images in real-time can be crucial which can be achieved by expanding the outcome of this study. There may be challenges in developing models that are both accurate and efficient enough for real-time analysis. Variability in imaging conditions (like lighting, resolution, and contrast) can affect model performance. Developing models robust to these variations is a significant challenge. Additionally, images may contain noise and artefacts, which can mislead the segmentation and clustering process. Developing methods to handle these efficiently is an ongoing challenge.

The objectives of some studies include classifying and quantifying cyanobacteria using deep learning techniques such as a Fast Regional Convolutional Neural Network (R-CNN) and convolutional neural networks. This suggests a move toward automated processes for segmentation, which can be more efficient and accurate than manual methods, depending on the quality and quantity of the training data. The SyMBac tool is an example of how researchers create synthetic images that closely resemble real micrographs, which can then be used to train accurate image segmentation models. If images involve segmentation based on synthetic training data, it would be an advanced application which require further development.

Combining imaging data with other omics data (like genomic or proteomic data) for a more comprehensive understanding of cyanobacteria and their harmful and/or nuisance metabolite is an area that is still under-explored. Integrating a portable automatic water sampling device with a microscopic imaging and photography system, complemented by the AI-based image analysis model, will significantly enhances access to different sampling sites. This synergy may not only minimize human error but also substantially enhances the ability to provide data promptly, achieving real-time or near real-time data delivery. Considering climate change impacts, effective integration of biological knowledge of cyanobacteria into the machine learning process, forecasting and modelling is a major knowledge gap (Hamilton et al, 2021). This integration is essential for more accurate and biologically relevant results.

The dominance of toxic and T&O compound-producing cyanobacteria is primarily a response to environmental stressors such as light, temperature, and nutrient availability. These stressors can also influence their morphology. However, the direct relationship between toxin and T&O compound production and changes in cell morphology is not straightforward and may vary among different cyanobacterial species. It is hypothesised that some cyanobacteria might exhibit noticeable morphological changes during variation in dominance of toxic strains and subsequent toxin and T&O production. For instance, Dapena et al. (2015) used imaging flow cytometry to observe changes in DNA content and nuclear and cell morphology occurring throughout clonal growth of toxic *Alexandrium minutum* Halim. Hence, integration of AI-based cell imagery to capture potential changes in cell morphology during changes in dominance of toxic strains requires further investigation.

Another important application is in enhancing image resolution. Techniques like super-resolution microscopy, which were once limited by the physical constraints of light diffraction, have been augmented by machine learning models that can predict high-resolution details from lower-resolution images. This has allowed researchers to view biological processes at a molecular level with unprecedented clarity.

## Conclusion

The results of this study demonstrated a promising approach in use of machine learning algorithms for segmentation and clustering cyanobacterial microscopic images, considering the varying morphologies and associated challenges (variable cell and colony density, presence of non-target objects, variable magnification, focal plane and lighting). The application of CNNs for the segmentation of cyanobacteria in microscopic images, as demonstrated in this study, marks a significant step forward in the field of digital image analysis and microbiology. While this study’s results are promising, future research should focus on comparing different CNN architectures, improving the robustness of the model against varying image qualities, and exploring its applicability to a wider range of microorganisms. This progression will not only enhance our understanding of microbial ecology but also pave the way for advancements in environmental monitoring and public health.

The study of cyanobacterial cell shapes through image clustering has significant practical applications. It aids in monitoring environmental health, particularly in detecting harmful algal blooms and on time implementation of mitigation strategies. It also contributes to our understanding of microbial ecology and the development of biotechnological applications. Further studies might focus on integrating more advanced machine learning techniques, such as deep learning, to enhance the accuracy of image clustering. This progress could unravel new aspects of cyanobacterial ecology and help in developing strategies to mitigate the impacts of algal blooms on water systems.

This study demonstrated that identifying knowledge gaps in machine learning-driven segmentation and shape clustering in cyanobacteria imaging requires understanding the intricacies of both cyanobacteria biology and advanced machine learning techniques. Segmentation accuracy and adaptability to variations, imaging conditions, dataset quality and availability, interpretability, and integration with biological knowledge require further thorough investigations.

## Acknowledgement

The authors acknowledge the support from water industry partners, and Monash University including Monash Department of Civil Engineering and Monash Faculty of Engineering academic establishment funds.

