## Supplementary Information for "Machine learning driven image segmentation and shape clustering of algal microscopic images obtained from various water types"

Table SI-1. Strengths and limitations of established referenced methods for studying algae and cyanobacteria*, and their application for research in ecology, physiology, and biotechnology (Lund et al., 1958; Weber, 1973; Wetzel and Likens 1991; Acker, 2003; American Public Health Association et al. 2017; Chorus and Welker, 2021; Vaughan et al., 2022).

| Light Microscopy | This is the most basic and widely used method. Light microscopy allows for the direct observation of cyanobacteria, enabling study of their morphology and behaviour. Although it offers limited resolution, it's essential for initial identification and basic research. |
| --- | --- |
| Electron Microscopy | This technique provides higher resolution images compared to light microscopy. Scanning Electron Microscopy (SEM) and Transmission Electron Microscopy (TEM) are commonly used to study the ultrastructure of cyanobacteria. SEM gives a detailed surface view, while TEM allows for the observation of internal structures. |
| Fluorescence Microscopy | Cyanobacteria can be studied using fluorescence microscopy, which involves staining the cells with fluorescent dyes. This method is useful for understanding the physiological state and metabolic activities of cyanobacteria. |
| Confocal Laser Scanning Microscopy (CLSM) | CLSM provides three-dimensional images of cyanobacteria, offering insights into their spatial structure. It's particularly useful for studying cyanobacteria in biofilms or within ecological samples. |
| Flow Cytometry | Though not a direct imaging method, flow cytometry is vital for quantifying and analyzing multiple characteristics of cyanobacteria, such as size, pigmentation, and internal complexity. |

* These microorganisms are renowned for their ability to perform oxygenic photosynthesis, similar to plants, thus significantly contributing to the production of oxygen in the atmosphere. This process was crucial in shaping the Earth's early atmosphere and enabled the evolution of complex life forms. In aquatic ecosystems, cyanobacteria are key primary producers, forming the base of the food web. They convert sunlight into energy, thus supporting a wide range of aquatic life. Additionally, certain cyanobacteria can fix atmospheric nitrogen, making it available to other organisms, thereby enriching nutrient-poor environments. The dual nature of cyanobacteria highlights their importance in sustaining life and the delicate balance required to maintain ecosystem health. Their role in global biogeochemical cycles, particularly in carbon and nitrogen cycling, underscores their significance in both terrestrial and aquatic environments (Chorus and Welker, 2021).

Table SI-2. Further examples of image segmentation to show the diversity of cells and flocs that the algorithm was able to successfully separate. The red outlines show examples of tracked cells within each image. This demonstrate the versatility of the process. Please note that below red cycles are only limited examples as it is difficult to show all the segmented shapes without covering the images with red circles.

| 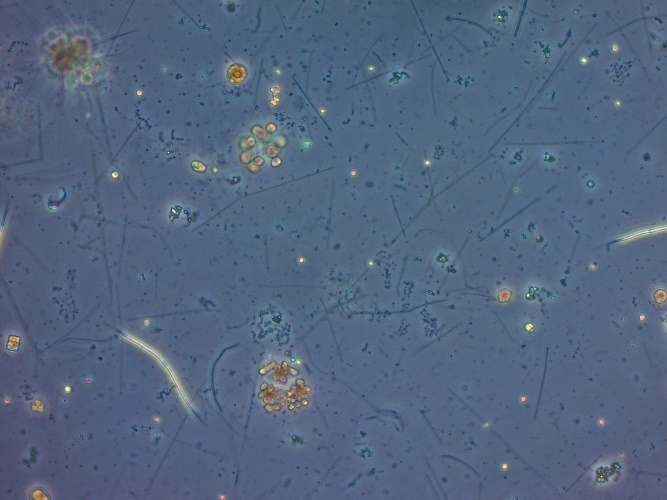 | 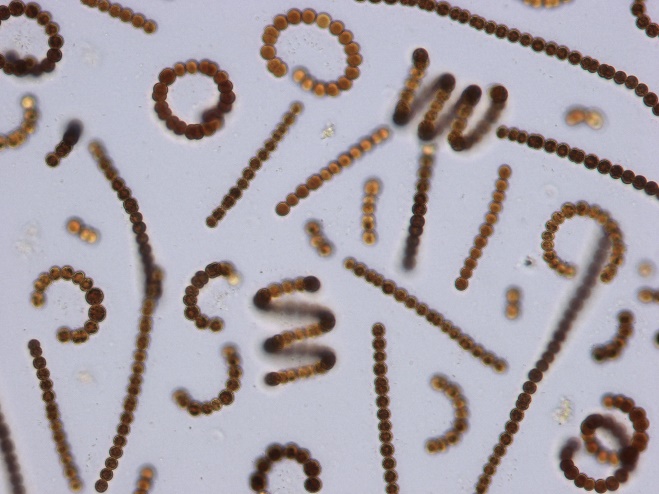 |
| --- | --- |
| 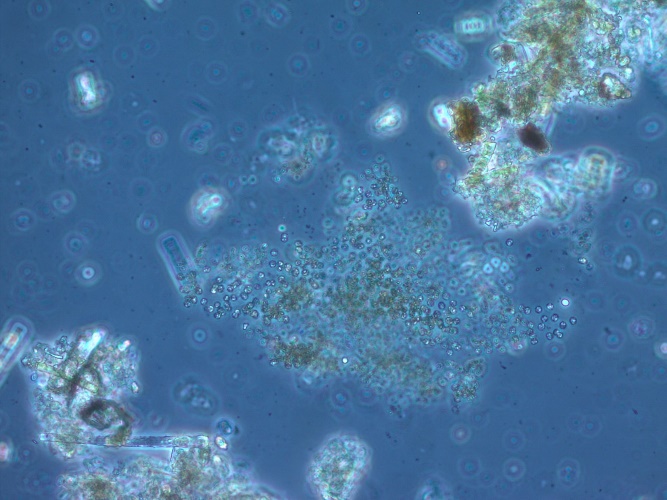 | 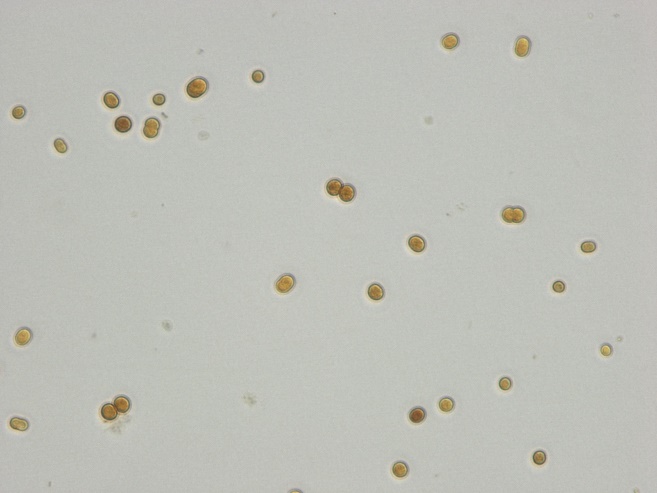 |
| 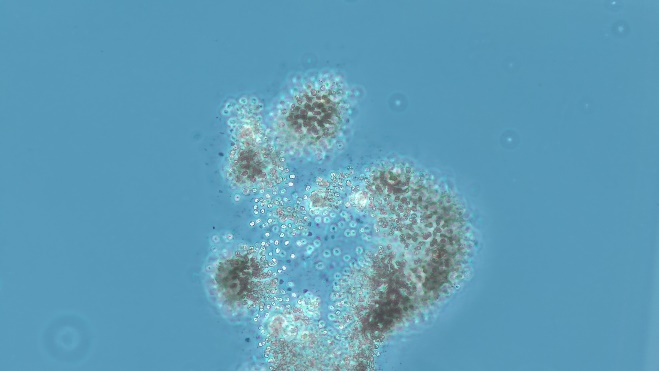 | 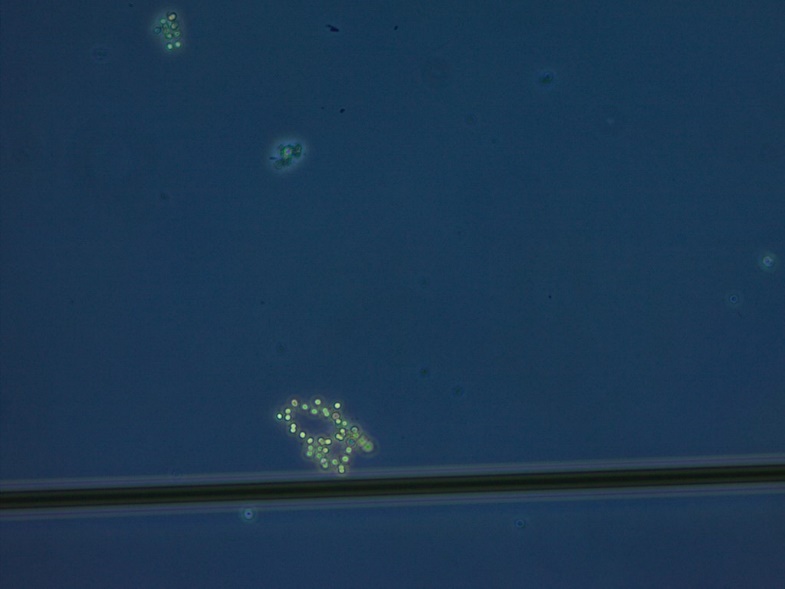 |
